# Non-destructive DNA extraction from specimens and environmental samples using DESS preservation solution for DNA barcoding

**DOI:** 10.1101/2024.08.09.607403

**Authors:** Eri Ogiso-Tanaka, Minako Abe Ito, Daisuke Shimada

## Abstract

DESS is a widely used storage solution for the preservation of DNA from biological tissue samples. DESS comprises 20% dimethyl sulfoxide, 250 mM ethylenediaminetetraacetic acid, and saturated sodium chloride, and its efficacy has been confirmed in various taxa and tissues. DESS enables the stable, long-term preservation of both sample morphology and DNA. However, to access the DNA, excising a portion of the sample was necessary. Although DNA is a valuable source of information for species identification, DNA extraction can result in the loss of an entire sample or segment, especially in small-sized organisms, thereby compromising specimen value. Therefore, establishing non-destructive DNA extraction techniques is imperative. Thus, this paper presents a protocol for conducting non-destructive DNA extraction and DNA barcoding using a portion of the DESS supernatant obtained from a nematode specimen. This method was successfully employed for DNA barcoding of nematodes that were stored in DESS at room temperature (-10∼35°C) for ten years. Moreover, the method can be potentially applied in the preservation and non-destructive extraction of DNA from specimens of various species. Following sample collection, a bulk environmental sample from sediment and seagrass is immediately immersed in DESS in the field. Subsequently, DNA is extracted from the supernatant solution, allowing non-destructive DNA barcoding. Overall, this paper presents comprehensive protocols for DNA extraction from DESS supernatants and demonstrates their practical application using meiofauna (small animals) and diatoms as examples.

## Introduction

DNA barcoding is a simple method for species identification based on DNA sequences unique to each species [1]. This approach is particularly beneficial for identifying taxa that are difficult to identify morphologically, such as free-living marine nematodes [2]. Considering that most free-living nematodes are small in body size (0.1–5.0 mm) and have no detachable appendages, the entire individuals are frequently consumed during DNA extraction and no voucher specimens remain. Although nematodes have been dissected to preserve the head and tail regions, which provide significant morphological information [3], and photographs have been taken before DNA extraction [4], sufficient information for taxonomic considerations is not always retained. In some cases, cryptic species have been revealed by analyzing sequences from specimens that were considered to be the same species based on morphology [5]. Therefore, sequenced specimens should be kept in an observable condition whenever possible. To address these issues, developing methods that enable DNA extraction from preserved small invertebrate specimens without any loss of morphological information is necessary.

Currently, preservation reagents, such as anhydrous ethanol and RNAlater Stabilization Solution (Thermo Fisher Scientific, USA), are employed to delay or inhibit DNA degradation during biological sample storage [6, 7]. However, these solutions cause tissue dehydration, which may affect morphology, and are not always optimal specimen preservation reagents for taxonomic purposes [8]. DESS is a saturated NaCl solution containing dimethyl sulfoxide (DMSO) and ethylenediaminetetraacetic acid (EDTA) and is widely used to preserve DNA in biological tissue samples. The solution was initially developed for preserving DNA in avian tissues at room temperature (R.T.) [9]. Subsequently, the solution was proven to preserve both physical structures and high molecular weight DNA in several marine invertebrates [10], including marine, terrestrial, and parasitic nematodes [4]. Furthermore, the maintenance of high molecular weight DNA by DESS at R.T. has recently been verified through molecular biology studies [11, 12].

DESS solution can also be employed as a preservation solution for environmental DNA samples. Soil has been stored in DESS to investigate bacterial and microbial community structures [13, 14] and fecal samples have been preserved in DESS to detect parasitic nematodes in dogs [15]. Withstanding these previous findings, we propose that environmental samples can be stored in DESS, and DNA can be easily extracted from the supernatant. Notably, DESS contains a high EDTA concentration and can dissolve calcium, which may result in the loss of morphology for some organisms that rely on calcium for their skeletal structure. Nevertheless, DESS is a broadly applicable preservation solution for other species and is capable of maintaining both morphology and DNA integrity, thereby facilitating non-destructive DNA extraction and DNA barcoding.

## Materials and Methods

The general sample preparation and DNA extraction protocols described in this peer-reviewed article are separately published on protocols.io (https://dx.doi.org/10.17504/xxxxxx) and are provided as files S1 and S2.

### Old nematode specimen samples

Free-living marine nematodes were collected from three localities on the Pacific coast of

Hokkaido, northern Japan (Table 1). Sandy sediments, seagrass roots from *Phyllospadix iwatensis* Makino (1931) and mussel byssuses from *Mytilus galloprovincialis* Lamarck (1819) or *Mytilus trossulus* Gould (1850) were collected by hand from the intertidal zone of rocky shores. In the laboratory, substrates were washed thrice in tap water and the supernatants were filtered through 63-μm mesh to separate meiofauna [16]. Nematodes were sorted under a stereomicroscope with an Irwin loop [17, 18] and preserved in 1.5 ml DESS (pH 8.0) [4]. DESS-preserved specimens (Figs 1 and 2A) were observed and identified using a differential interference contrast microscope (Olympus BX51).

**Fig 1.**
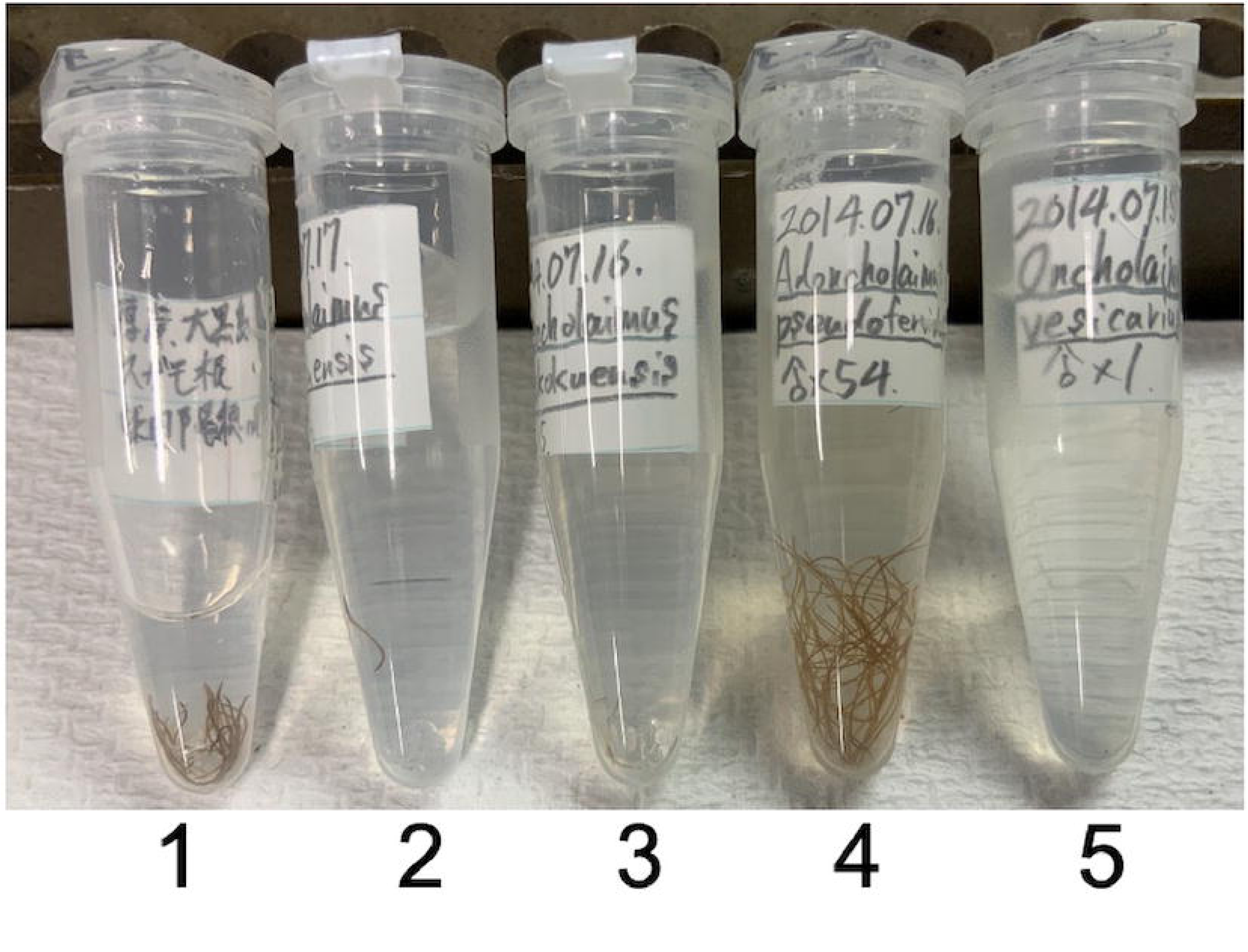
Ten-year-old DESS preserved specimens used in this study. The supernatant for Sample 1 (the leftmost) was already used for DNA extraction. The number indicates the sample number.

**Fig 2.**
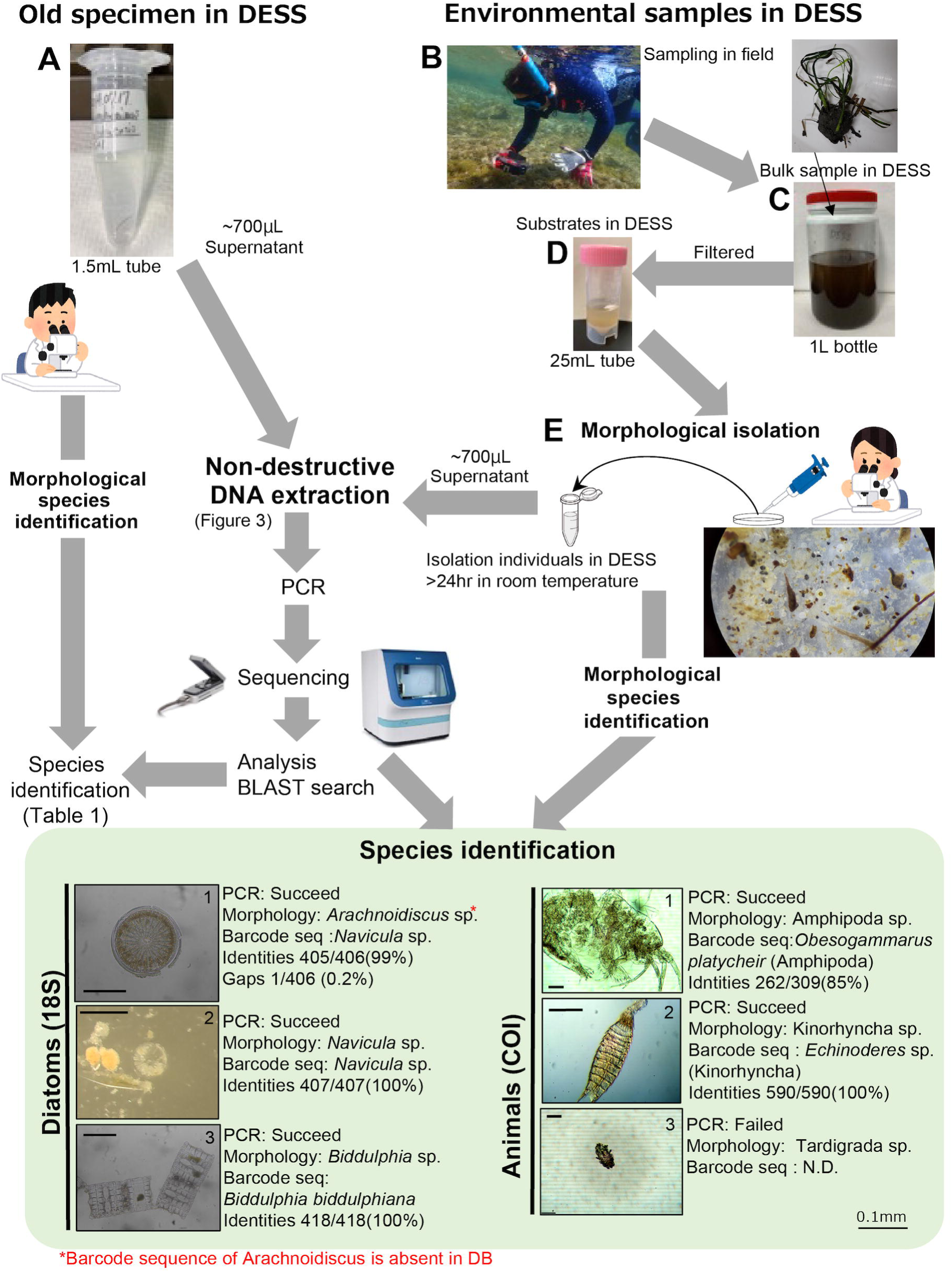
Workflow and practical examples of sample preparation, DNA barcoding, and species identification results. Old specimen in DESS: the workflow of old nematode specimen DNA barcoding. **A.** Old nematode specimen stored for ten years at room temperature (-10∼35°C) in DESS. Environmental samples in DESS: the workflow involves transporting environmental samples immersed in DESS to the laboratory, where small organisms are isolated for DNA barcoding and morphological observation. B. Sampling procedure in the field. C. One-liter bottle containing the entire seagrass (with sandy rhizomes), sand, and small organisms, in DESS. D. Small organisms separated by filtering C through a 63 μm mesh filter (see Materials and Methods). E. Microscopic observation and isolated individuals.

**Table 1.** Information on the nematodes examined in this study.

| Tube number | Species | n* | Body length | Collection date | Locality | Lat./Lon. | Habitat |
| --- | --- | --- | --- | --- | --- | --- | --- |
| 1 | *Adoncholaimus daikokuensis*<br>Shimada & Kajihara, 2014 | 21 | ca. 5 mm | 15 Jul. 2014 | Daikoku Island,<br>Hokkaido, Japan | 42°57'41"N<br>144°52'25"E | among *Phyllospadix*<br>roots |
| 2 | *Adoncholaimus daikokuensis* | 1 | ca. 5 mm | 17 Jul. 2014 | Shinhidaka,<br>Hokkaido, Japan | 42°17'03"N<br>142°27'47"E | among *Phyllospadix*<br>roots |
| 3 | *Adoncholaimus daikokuensis* | 5 | ca. 5 mm | 16 Jul. 2014 | Hiroo, Hokkaido,<br>Japan | 42°14'01"N<br>143°18'58"E | among *Phyllospadix*<br>roots |
| 4 | *Adoncholaimus pseudofervidus*<br>Shimada & Kajihara, 2014 | 54 | ca. 10 mm | 16 Jul. 2014 | Hiroo, Hokkaido,<br>Japan | 42°14'01"N<br>143°18'58"E | among *Mytilus*<br>byssuses |
| 5 | *Oncholaimus* sp. | 1 | ca. 5 mm | 15 Jul. 2014 | Daikoku Island,<br>Hokkaido, Japan | 42°57'41"N<br>144°52'25"E | fine sand |

\* n represent the number of individuals present in the tube.

Environmental samples in DESS: the workflow involves transporting environmental samples immersed in DESS to the laboratory, where small organisms are isolated for DNA barcoding and morphological observation. **B.** Sampling procedure in the field. **C.** One-liter bottle containing the entire seagrass (with sandy rhizomes), sand, and small organisms in DESS. **D.** Small organisms separated by filtering C through a 63-μm mesh filter (see Materials and Methods). **E.** Microscopic observation and isolated individuals.

### Environmental samples containing various species

Environmental samples were collected from two localities in southern Chiba, Japan; Emi (35°3′52″N, 140°4′55″E) and Chikura (34°56′36″N, 139°57′51″E) on July 12, 2022. The samples were collected by uprooting seagrass shoots from *Phyllospadix japonicus* Makino (1897) from intertidal rocks (Fig 2B). The roots were detached by pulling the bottom of the shoot by hand, and a metal scraper was used if the roots were firmly attached. To minimize the loss of sediment and associated organisms, detached *P. japonicus* was carefully placed in a plastic bag and brought to the beach. At the beach, *P. japonicus* shoots with sediments were transferred to a plastic container filled with DESS and transported to the laboratory (Fig 2C). No temperature control was performed during the entire procedure.

In the laboratory, *P. japonicus* shoots with sediments and organisms in DESS were stirred in tap water to remove lighter components, including living organisms, and the supernatants were filtered through 63-μm mesh to separate organisms (Fig 2D, S1 File). Meiofauna and diatoms were sorted under a stereomicroscope (Fig 2E) and preserved in separate 2.0-ml tubes with DESS. They were then identified using a differential interference contrast microscope (Nikon OPTIPHOT).

### DNA extraction from DESS solution supernatant

All subsequent steps were performed using sterile filter tips and low DNA binding tubes (e.g. Eppendorf tube, Cat.#0030108051). A 10-µl aliquot of silica solution (wakosil; FUJIFILM, Wako Pure Chemical Corporation Japan, Cat.#232-00841) with 0.01 N HCl (2.5 g wakosil in 11 ml of 0.01 N HCl) was added to a new 1.5-ml tube (b). The supernatant (500 µl) was then transferred from the DESS solution containing the stored target organisms into the 1.5-ml tube. An equal volume (500 µl) of binding buffer (5 M guanidine thiocyanate, 4% Triton X-100, 10% Tween 20, 10 mM sodium acetate) was added and mixed thoroughly. An equal volume (500 µl) of ethanol was then added and mixed well. When the liquid volume was substantial, the liquid was divided among multiple tubes. After mixing, the samples were vortexed and incubated at R.T. for 5 min. The tubes were then centrifuged at 10,000 rpm (9,100 xg) for 1 min. The liquid phase was then discarded by pipetting or using an aspirator, and the pellet was washed twice with 500 µl of wash buffer prepared by diluting a stock solution (10 mM TrisHCl [pH 8.0], 100 mM NaCl, 0.5 mM EDTA [pH 8.0]) with ethanol at a 1:4 ratio (stock solution:ethanol). The pellet was air dried after washing. TE buffer (203 µl) was added to the dried silica pellet and vortexed. The pellet was then incubated at R.T. for 5 min to release DNA from the silica particles and centrifuged at 10,000 rpm for 1 min. The supernatant (200 µl) was transferred to a new tube containing 1/10 volume (20 µl) 3 M sodium acetate (pH 5.2) and 1 µl glycogen. Equal volumes (200 µl) of isopropanol was added, mixed well, and centrifuged at 15,000 rpm (20,400 xg) for 30 min at 4 °C. The liquid phase was discarded using a pipette or aspirator, and the pellet was washed twice with 500 µl of 75% ethanol. The liquid phase was completely discarded using a pipette or aspirator. The pellet was air dried for 30 min under dark conditions; in the absence of glycogen, the pellet may not be visible. The pellet was resuspended in 10 µl of 0.1× TE buffer by tapping, and the DNA solution was incubated for 10 min at R.T. Given that the DNA concentration was below the limit that could be measured by NanoDrop (Thermo Fisher Scientific, USA), measurement was unnecessary. The 1 μl of DNA was used for polymerase chain reaction (PCR) without adjusting the DNA concentration.

### PCR for DNA barcoding

PCR was performed using indexed universal primers (Table 2, S1 Table). PCR mixtures (20 μl final volume) contained 1 μl of DNA, 1 μl of each indexed primer (2 μM), 10 μl of 2× Gflex Buffer, 0.2 μl of Tks Gflex DNA Polymerase, and 6.8 μl of ddH2O. Cycling conditions for COI (mlCOIinf/HCO2198) for animals except for nematodes were as follows: one cycle at 94 °C for 2 min, followed by 40 cycles at 94 °C for 30 s, 48 °C for 30 s, and 72 °C for 1 min, and final extension at 72 °C for 1 min [19, 20]. Cycling conditions for COI (JB3/JB5) for nematodes were as follows: once cycle at 94 °C for 2 min, followed by 40 cycles at 94 °C for 30 s, 48 °C for 30 s, and 72 °C for 30 s, and final extension at 72 °C for 1 min [21, 22]. Cycling conditions for 18S (D512/D978) for diatoms were as follows: one cycle at 94 °C for 2 min, followed by 40 cycles at 94 °C for 30 sec, 49 °C for 30 s, and 72 °C for 30 s, and final extension at 72 °C for 1 min [23]. Amplified DNA fragments were checked using agarose gel electrophoresis or Agilent 4200 TapeStation D1000 ScreenTape (Agilent Technologies, Inc., USA). When PCR product amount was very low, purification was performed before repeating PCR. For purification, 1.5× Serapure (S2 File) or AMpure (Beckman Coulter, USA) beads were added to the first PCR products to remove primers and salt. The mixture was then mixed well and allowed to stand for 5 min. The tube was placed on a magnetic stand, and the supernatant was discarded when it became clear. The beads were washed twice with 75% ethanol, and all the supernatant was carefully removed. The beads were then air dried, ensuring not to over dry. The 0.1xTE buffer (6 μl) was added and mixed well. The tube was then placed on a magnetic stand, and 1 μl of the supernatant was used to perform PCR again under the same conditions. The composition of Serapure is as follows: 2% SpeedBeads Carboxylate-Modified Magnetic Particlescmmp (Sigma-Aldrich, USA), 18% PEG8000, 1 M NaCl, 10 mM Tris-HCl [pH 8.0] 1 mM EDTA [pH 8.0], and 0.05% Tween 20 [24] (S2 File).

**Table 2.**
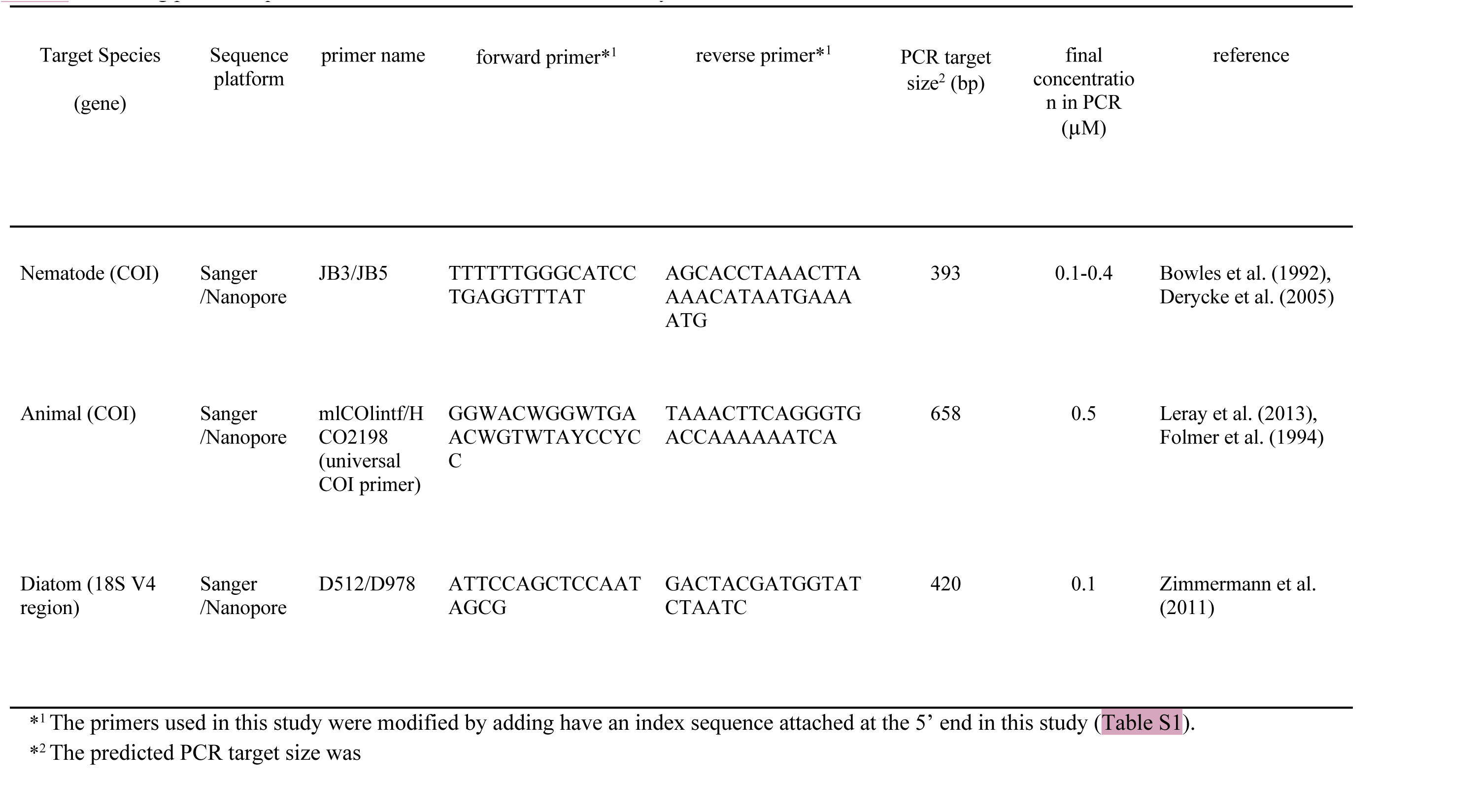
Barcoding primer sequences and concentrations used in this study.

### Library preparation and sequencing

#### Sanger platform

PCR products from nematode specimens were purified using a NucleoSpin Gel and PCR Clean-up kit (MACHEREY-NAGEL, Germany). Nucleotide sequences were determined by direct sequencing using a BigDye Terminator Kit ver. 3.1 (Applied Biosystems, USA) with a 3500xL Genetic Analyzer (Applied Biosystems). The JB3/JB5 primer [21, 22] sets were the same as those used for PCR. DNA sequences were aligned using MEGA X [20].

#### Nanopore platform

PCR products were mixed in equal amounts, purified using Serapure beads at a 0.8× ratio, and dissolved in EB buffer. The DNA concentration was measured for the purified PCR products using Qubit high sensitivity (Thermo Fisher Scientific). The purified 500 ng of PCR products were mixed with 3.5 μl of Ultra II End-prep buffer, 1.5 μl of UltraII End-prep enzyme (Cat No. E7546S, New England Biolabs, USA), and dH2O up to 30 μl in a 0.2-ml PCR strip. The mixture was then inculcated at 20°C for 30 min for the end-repair reaction, followed by 65°C for 5 min for the dA-tailing reaction. The mixture was then purified using an equal volume (30 μl) of AMpure or Serapure, according to the manufacturer’s instructions, using Nanopore ligation kit protocols (Oxford Nanopore Technologies, Oxford, UK). After washing the beads, 25 μl of dH2O was added to elute dA-tailed DNA and was mixed by pipetting. The mixture was then left to stand for 2 min. The 24 μl of dA-tailed DNA and 2 μl of ligation adapter were ligated using 4 μl of Quick T4 DNA Ligase (New England Biolabs) in 40 μl of reaction volume. After ligation, the mixture was purified using 20 μl of AMpure or Serapure beads and washed twice with 125 μl of Short Fragment Buffer. After washing beads, 7 μl of EB buffer was added to elute the DNA library. The mixture was mixed by pipetting and left to stand for 1 min. Before sequencing, the concentration was confirmed using 1 μl of DNA library with the Qubit High Sensitivity kit (Thermo Fisher Scientific). Subsequently, 15 μl of Sequencing Buffer, 10 μl of Library Beads, and 5 μl of DNA library were mixed. Sequencing was performed using Flongle (R9.4) flow cell on a MinION Mk1b portable sequencer (Oxford Nanopore Technologies).

### Sequencing analysis for Nanopore read

The MinKNOW run using FLO-FLG001 flow cell and the SQK-LSK109 sequencing kit generated FAST5 files in the “pass” folder, which were then converted to FASTQ files using Guppy ver. 6 (Oxford Nanopore Technologies). Subsequently, barcode sequences were detected using two analysis tools. First, ONTbarcoder (https://github.com/asrivathsan/ONTbarcoder) was used for demultiplexing and consensus sequence calling from FASTQ files [25]. ONTbarcoder was a good tool for DNA barcoding of a single species, especially for amino acid sequences (such as COI barcoding of animals), with the translation-based error correction; however, the tool was not effective when contaminant reads were mixed. Second, NGSpeciesID (https://github.com/ksahlin/NGSpeciesID) was used for consensus sequence calling from demultiplexed files [26]. NGSpeciesID was also a good tool for DNA barcoding of samples containing a small number of mixed species; however, this tool did not have an error correction function.

### BLAST search and comparison of Sanger and Nanopore sequencing

BLASTN search (https://blast.ncbi.nlm.nih.gov/) was used to identify the species from the sequences. DNA sequences were aligned using muscle in MEGA X [27] and Parasail [28] in SnapGene (https://www.snapgene.com/).

## Results and Discussion

### DNA barcoding of old specimen samples using non-destructively extracted DNA

Using the protocols described (Figs 2, 3), we were able to reliably obtain DNA, albeit at very low concentrations. The extracted DNA concentration was below the detection limit for both NanoDrop (Thermo Scientific) and Qubit (Thermo Fisher Scientific). However, PCR with the JB3/JB5 primer (S1 Table) amplified the COI region in all DNA samples (Fig 4). For the low-amplification sample (Sample 5 in Fig 4A), the first PCR product was purified using magnetic beads and then used as a template for the second PCR using the same JB3/JB5 primer set (Fig 4B). Using these PCR products, Sanger sequencing was performed (Fig 5) and barcode sequences were obtained. DNA libraries for the Nanopore platform were constructed from the same PCR products and DNA barcoding was also performed with Nanopore Flongle flow cell (ONT). As a result of barcoding using Nanopore, multiple species were detected from Sample 3 and 5 (Table 3). For Sample 3, DNA barcoding using ONTbarcoder [25] only yielded the consensus sequence of *Adoncholaimus daikokuensis*. However, when the same data was analyzed using NGSpeciesID [26], two nematode species were detected: *A. daikokuensis* (475 out of 606 reads) and *Litinium* sp. (98 reads). In Sanger sequencing, low background noise was observed in the sequencing chromatograms, and the sequence of *A. daikokuensis* was obtained (Fig. 5).

**Fig 3.**
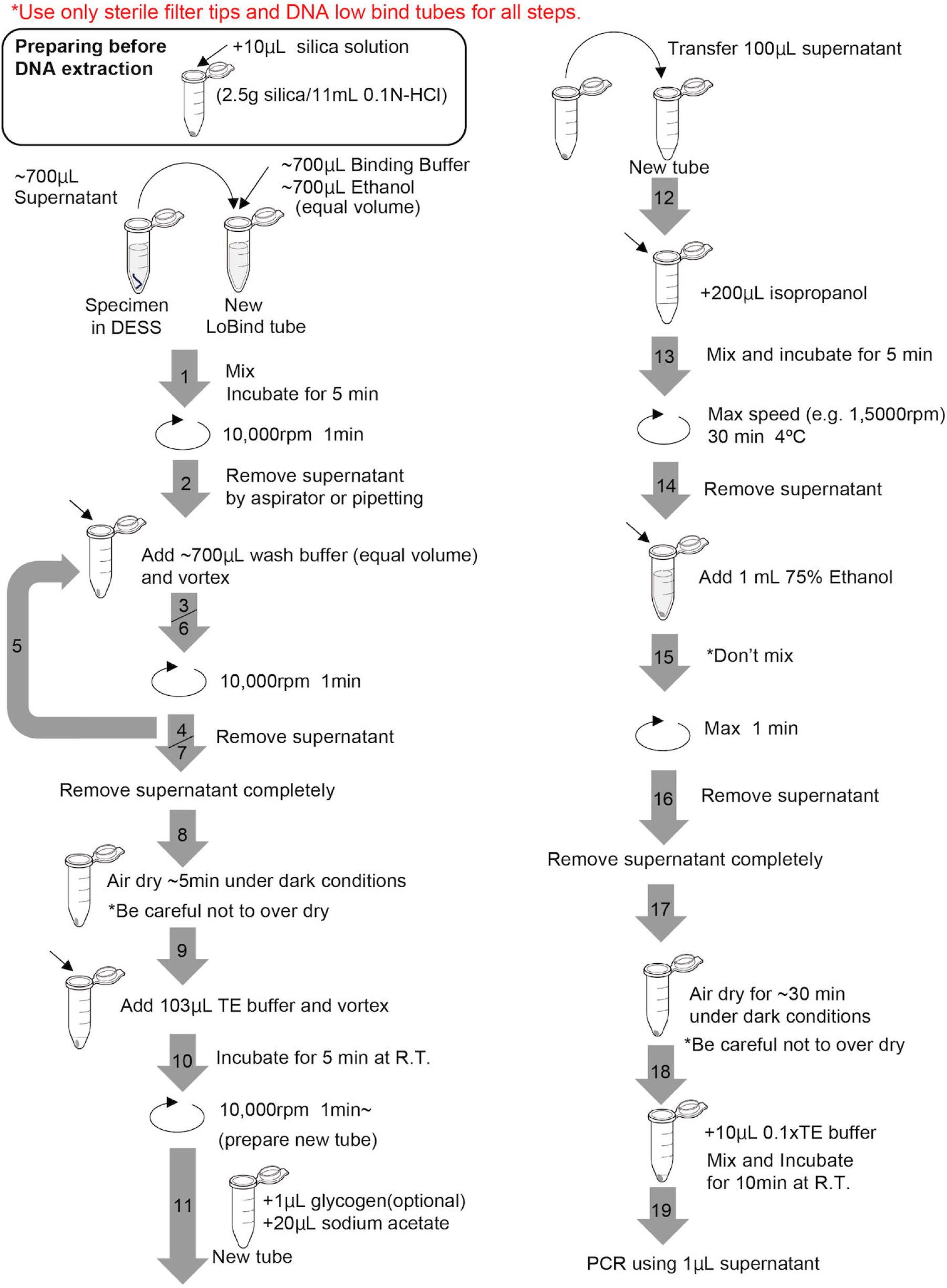
Schematic diagram of non-destructive DNA extraction protocol.

**Fig 4.**
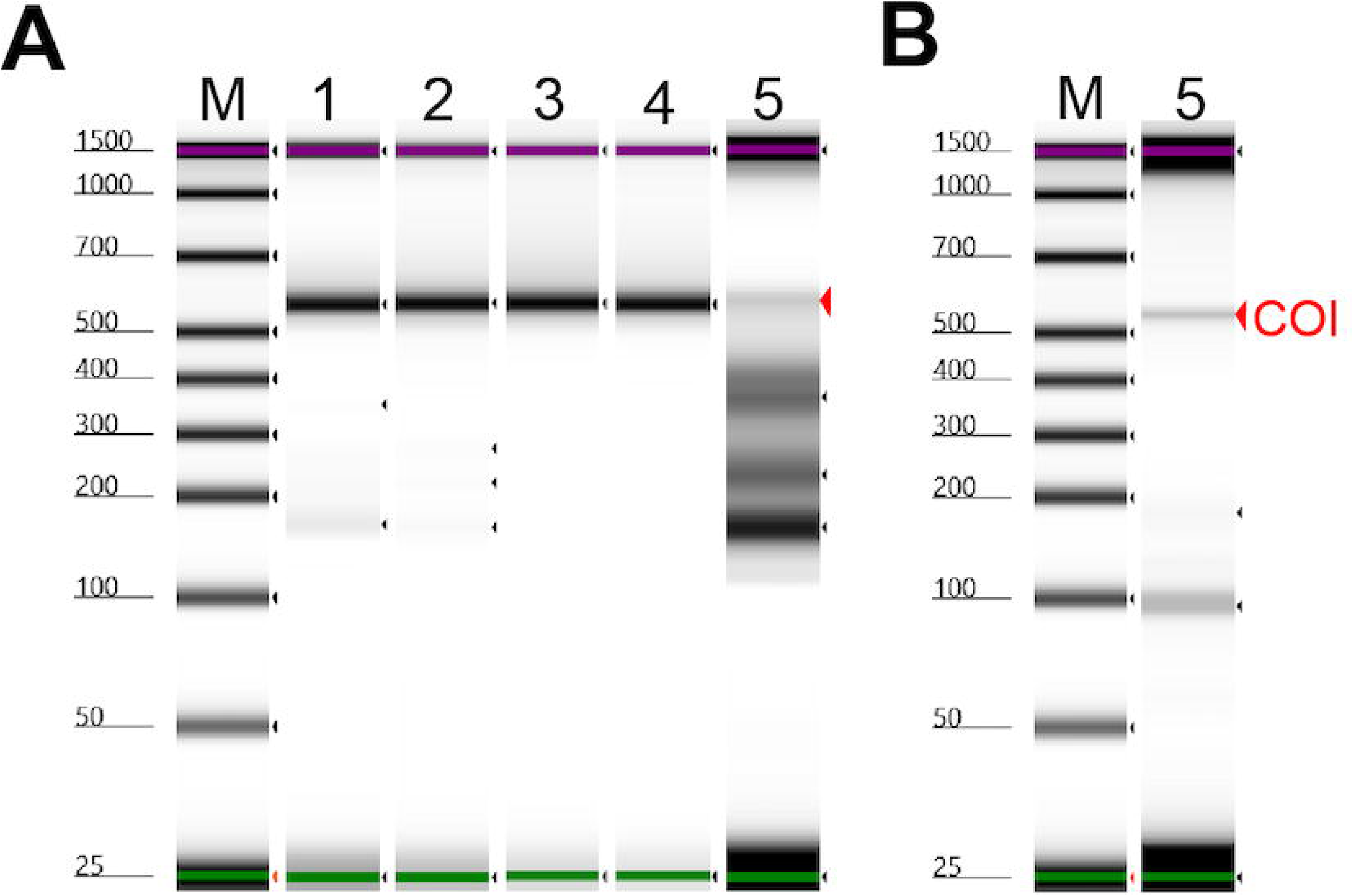
Polymerase chain reaction (PCR) products amplified using JB3/JB5 primer sets with DNA extracted non-destructively from nematode specimens stored for 10 years. **A.** Samples 1–4 exhibited single bands during amplification, whereas Sample 5 did not yield sufficient PCR amplification. **B.** PCR products from the second round of PCR, where the first PCR product of Sample 5 was purified and used as a template with the same primer set. M represents the size marker, and 1–5 denote sample numbers.

**Fig 5.**
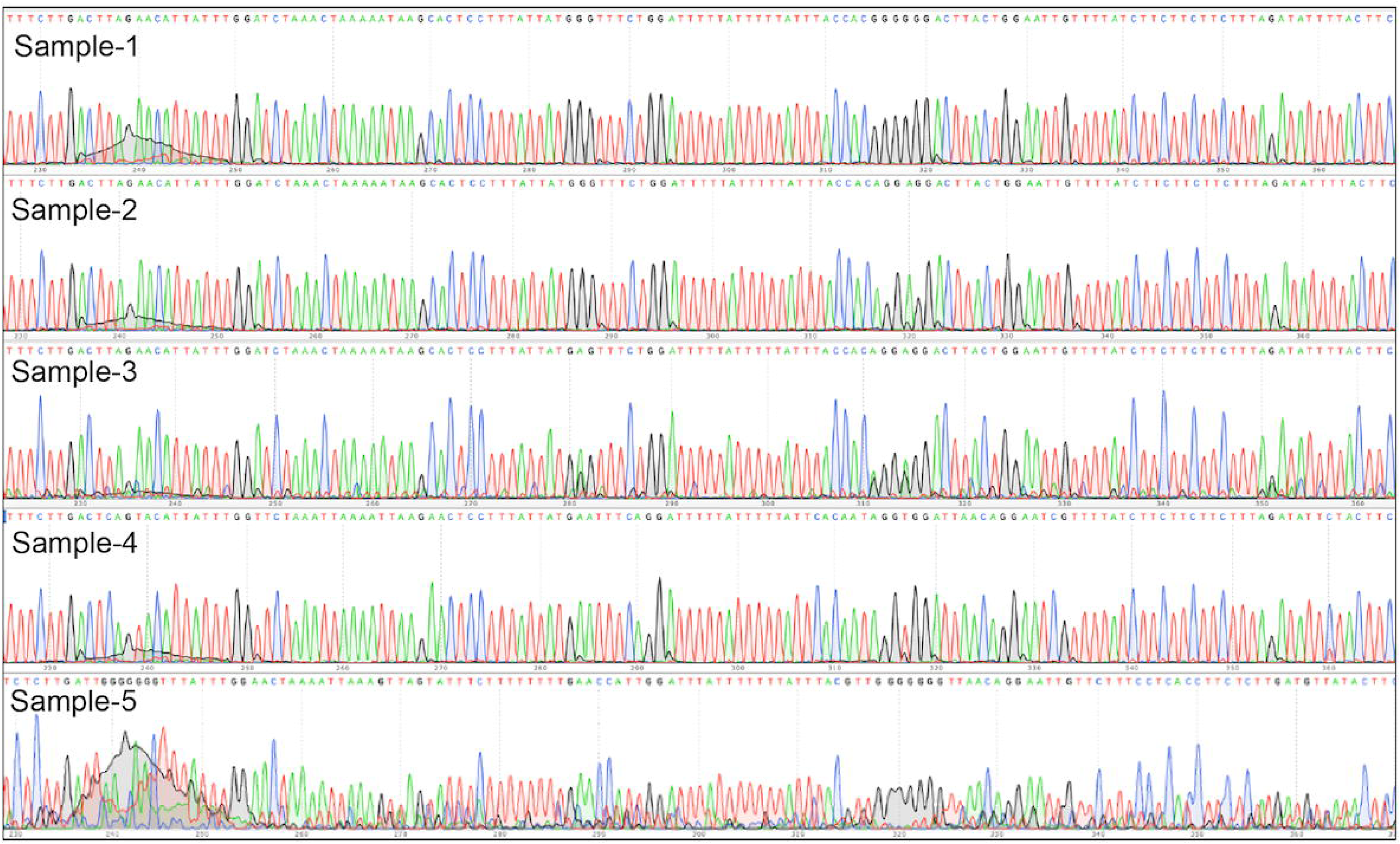
Sangar sequence chromatograms of sample 1–5.

**Table 3.**
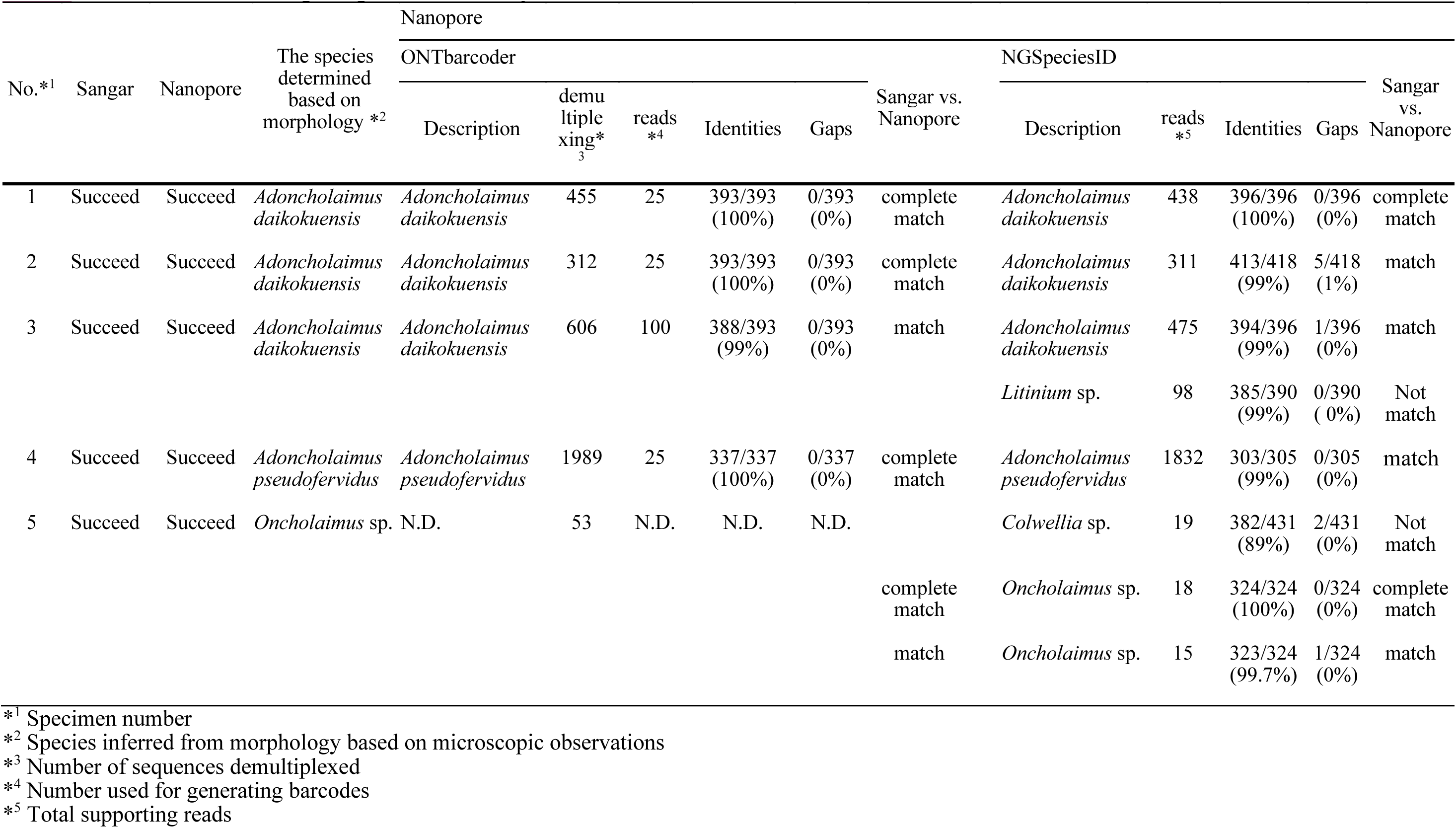
Result of DNA barcoding using non-destructively extracted DNA.

Considering that majority of reads (80%) were from *A. daikokuensis*, even if DNA from multiple species was mixed, the analysis by Sanger sequencer or ONTbarcoder with Nanopore would not be affected. However, we were not familiar with *Litinium* sp., and considering the low possibility of it being a contaminant, the sequence was highly likely to be obtained from the stomach contents. This result suggested that by using next-generation sequencing (NGS) and tools that output multiple consensus sequences, such as NGSpeciesID [26], more detailed information can be obtained from specimens. For Sample 5, no consensus sequence could be generated using ONTbarcoder [25]. When analyzed with NGSpeciesID [26], three output sequences were generated, with the first sequence identified as *Colwellia* sp. (Bacteria) and the second and third were two consensus sequences of *Oncholaimus* sp. with a difference of one base and one gap (S1 Fig.). Although the background noise during Sanger sequencing was higher than that of the other four samples, the sequence for *Oncholaimus* sp. could be obtained (Fig 5). Furthermore, although it was difficult to read several tens of bases from the primer region during Sanger sequencing, using NGS enabled us to obtain sequences immediately following the primer region, resulting in the acquisitions of barcode region sequences 60–70 bp longer than those obtained using Sanger sequencing (S1 Fig.). For Sample 5, which did not show sufficient amplification after one round of PCR, only 53 reads could be obtained. Due to the lower PCR product amount compared with that of other samples, good results could not be achieved. Additionally, the bacterial *Colwellia* sp. accounted for 36% of the reads; hence, no consensus sequence could be obtained with ONTbarcoder. Therefore, DNA barcoding is possible with Sanger sequencing as long as there are approximately 63% reads of one species, even if other species are mixed in. Based on these results, selecting the appropriate sequencing platform and analysis tool according to the objective is necessary. Furthermore, we used the R9.4.1 kit and Flongle flow cell for the Nanopore platform; however, we recommend using the latest reagents within their expiration dates to improve accuracy.

### Environmental samples containing various species

Using the protocols described (Figs 2B-E and 3), we obtained DNA from DESS supernatant containing sorted organisms (Fig 2E). The extracted DNA concentrations were below the detection limits for both NanoDrop (Thermo Scientific) and Qubit (Thermo Fisher Scientific).

After 24 h, we extracted DNA using 500 µl of DESS supernatant out of 2.0 ml (Fig 2E). PCR amplification using universal COI primers (mlCOIinf/HCO2198) for meiofauna [19,20] and 18S barcoding primers (D512/D978) for diatoms [23], yielded the desired PCR products in five out of six samples (Fig 2). For Animal-3, whose COI failed to amplify during PCR, the DNA may have degraded owing to the organism already being dead or the primer sequences may have mismatched. DNA libraries were constructed from these five PCR products and DNA barcoding was performed with Nanopore Flongle (ONT). As a result, identification based on morphology and barcode sequence matched for four out of five samples (Table 4, S3 Table). Animal-1 was morphologically identified to be Amphipoda sp., whereas no consensus sequence was obtained with ONTbarcoder [25]. When using NGSpeciesID [26], two amphipod species sequences were obtained, with 21 reads for the first species and 92 reads for the second species. The similarity between these two consensus sequences was as low as 77%; hence, a sequencing error was unlikely (S2 Fig). A single individual was isolated under the microscope, but contamination may have occurred. To confirm that the two COI sequences were from individual species, their corresponding amino acid sequences were checked. When the nucleotide sequences were translated into amino acids, the second consensus sequence could be translated into amino acids over the entire coding region, whereas two stop codons were found in the first consensus sequence, even after considering sequencing errors (S2 Fig). This strongly suggests that the COI was duplicated in the Animal-1 sample, with one copy being a pseudogene. Sanger sequencing revealed that the obtained sequence matched the second consensus sequence derived from NGSpeciesID (S3 Fig). Upon closer inspection of the chromatogram, minor peaks that seemingly corresponded with the first consensus sequence were observed, although they did not affect the overall result (S3 Fig). Amplification was successful for Animal-2, and 3275 reads were obtained, with most of the reads on a single barcode sequence. The obtained sequence was that of Kinorhyncha sp. and was consistent with the identification by morphological observation. For such samples, successful barcoding results can be obtained regardless of the sequencing platform or analysis tool. For diatom samples, successful PCR amplification was conducted for all samples, and each sample yielded a sufficient number of reads, exceeding 1000. Given that 18S was used for the analysis, the translation-based error correction of ONTbarcoder could not be utilized, and NGSpeciesID yielded slightly better results. Diatom-1 was identified as *Arachnoidiscus* sp. based on its morphology; however, because no sequence data for *Arachnoidiscus* species exists in the NCBI database, the accuracy of the consensus sequence could not be determined (Table 4). Diatom-2 was morphologically identified as *Navicula* sp., the barcode sequence matched 99% with *Navicula* sp. when analyzed with ONTbarcoder [25] and matched 100% with NGSpeciesID [26]. For Diatom-3, the barcode sequence also matched 100% with *Biddulphia* sp. using both tools and with the morphological identification (Table 4). However, with NGSpeciesID, in addition to the 567 reads of the *Biddulphia biddulphiana* sequence, 264 reads of the polychaete *Neoamphitrite figulus* (Annelida) considered a contaminant, were also detected (Table 4). ONTbarcoder [25] correctly obtained the consensus sequence of *B. biddulphiana*, even when mixed with other species. This is most likely because the total enough reads and contaminating species were low similarity sequence. In cases where similar sequences (duplicate genes or closely related species) are mixed, as was the case with Animal-1, NGSpeciesID needs to be used instead of ONTbarcoder.

**Table 4.** Result of DNA barcoding using non-destructively extracted DNA from isolated microorganisms.

| Sample No. | Number of individuals / tube | PCR | The species determined based on morphology*<sup>1</sup> | Nanopore |  |  |  |  | Morphology vs. barcode seq | NGSpeciesID |  |  |  | Morphology vs. barcode seq |
| --- | --- | --- | --- | --- | --- | --- | --- | --- | --- | --- | --- | --- | --- | --- |
|  |  |  |  | ONTbarcoder*<sup>2</sup> |  |  |  |  |  |  |  |  |  |  |
|  |  |  |  | Description | demultiplexing*<sup>3</sup> | reads*<sup>4</sup> | Identities | Gaps |  | Description | reads*<sup>5</sup> | Identities | Gaps |  |
| Animal-1 | 1 | Succeed | Amphipoda sp. | N.D. | 143 | N.D. | N.D. | N.D. | N.D. | *Obesogammarus platycheir* (Amphipoda) | 21 | 262/309 (85%) | 1/309 (0%) | match (Amphipoda) |
|  |  |  |  |  |  |  |  |  |  | Ischyroceridae sp. (Amphipoda) | 92 | 272/313 (87%) | 0/313 (0%) | match (Amphipoda) |
| Animal-2 | 1 | Succeed | Kinorhyncha sp. | *Echinoderes* sp. (Kinorhyncha) | 3275 | 25 | 590/590 (100%) | 0/590 (0%) | completely match | *Echinoderes* sp. (Kinorhyncha) | 3272 | 590/590 (100%) | 0/590 (0%) | completely match |
| Animal-3 | 1 | failed | Tardigrada sp. | - |  |  | - | - | - | - | - | - | - | - |
| Diatom-1 | 18 | Succeed | *Arachnoidiscus* sp. | *Navicula* sp. | 2557 | 100 | 399/401 (99%) | 0/401 (0%) | Not match | *Navicula* sp. | 473 | 405/406 (99%) | 1/406 (0%) | Not match |
|  |  |  |  |  |  |  |  |  | No sequence of *Arachnoidiscus* in NCBI db. |  |  |  |  | No sequence of *Arachnoidiscus* in NCBI db. |
| Diatom-2 | 17 | Succeed | *Navicula* sp. | *Navicula* sp. | 2086 | 100 | 400/401 (99%) | 0/401 (0%) | match | *Navicula* sp. | 428 | 407/407 (100%) | 0/407 (0%) | completely match |
| Diatom-3 | 20 | Succeed | *Biddulphia* sp. | *Biddulphia biddulphiana* | 1911 | 100 | 404/404 (100%) | 0/404 (0%) | completely match | *Biddulphia biddulphiana* | 567 | 418/418 (100%) | 0/418 (0%) | completely match |
|  |  |  |  |  |  |  |  |  |  | *Neoamphitrite figulus* | 264 | 405/405 (100%) | 0/405 (0%) | contamination (animal) |

\*<sup>1</sup> Refer to Figure 2 for microscopic images of each microorganism. \*<sup>2</sup> The barcode sequence of diatom generated by ONTbarcoder was result of Consensus by similarity. \*<sup>3</sup> Number of sequences demultiplexed \*<sup>4</sup> Number used for generating barcodes in animals, or coverage for closest sequences(90% cutoff) in diatom. \*<sup>5</sup> Total supporting reads

## Conclusion

DESS solutions preserve both the morphology and DNA of small algae and animals. Furthermore, DESS solutions have the potential to non-destructively extract DNA from these organisms for DNA barcoding. In this study, DNA barcoding was successfully performed with both individual specimens and environmental bulk samples using both Sanger sequencing and NGS. Moreover, using NGS may be appropriate to detect DNA dissolved from the gut contents of target specimens and other associated organisms.

## Data Availability

Sequencing data generated with this protocol are publicly available on the Sequence Read Archive (SRA, NCBI) under BioProject PRJDB16817.

## Acknowledgments

Authors thank Dr. Makoto Manabe and Dr. Utsugi Jinbo (National Museum of Nature and Science) for their support. Authors are also grateful to Dr. Akito Ogawa and Dr. Genki Kobayashi (National Museum of Nature and Science) for their constructive comments on the previous versions of the manuscript.

## Supplementally figure legend

**Figure S1** Sequence alignment showing the sequence similarity between consensus sequence 2 and 3 in specimen-5 (Table 4). Yellow boxed regions indicate the sequence registered in NCBI.

**Figure S2** A Sequence alignment showing the sequence similarity between consensus sequence 1 (best hit: Obesogammarus platycheir) and 2 (best hit: Ischyroceridae sp.) in Amimal-1 (Table 5). The sequence exhibit about 77% similarity (identity 250/324, gap 8/324). Red characters represent mismatch alignment. B,C Consensus sequence 1 (B) and 2 (C). The amino acid sequence was predicted. The asterisk (*) with a red background denotes the stop codon.

**Figure S3** Sanger sequencing chromatograms of both forward and reverse strands for the COI region (mlCOIintf/HCO2198) in Animal-1 (Table 5). Sequence alignment showing the sequence similarity among Sanger and NGS outputs [consensus sequence 1 (NGS-1: blast hit, Obesogammarus platycheir) and 2 (NGS-2: blast hit, Ischyroceridae sp.) in Table S3].

## Supporting information

**S1 File. Protocol for nematode sample preparation (Protocols.io). S2 File. Protocol for non-destructive DNA extraction (Protocols.io).**

**S1 Fig. Sequence alignment between consensus sequences 2 and 3 in specimen 5.** Yellow boxed regions indicate the sequence registered in NCBI.

**S2 Fig. Sequence alignment and amino acid sequences of consensus sequences in Animal-1. A.** Sequence alignment showing the sequence similarity between consensus sequence 1 (best hit: *Obesogammarus platycheir*) and 2 (best hit: Ischyroceridae sp.) in Animal-1 (Table 5). The sequence exhibited approximately 77% similarity (identity 250/324, gap 8/324). Red characters represent mismatch alignment. **B.** Consensus sequence 1. **C.** Consensus sequence 2. The amino acid sequence was predicted. The asterisk (*) with a red background denotes the stop codon.

**S3 Fig. Sanger sequencing chromatograms of both forward and reverse strands for the COI region in Animal-1.** Sequence alignment showing the sequence similarity among Sanger and NGS outputs. Consensus sequence 1 (NGS-1: blast hit, *Obesogammarus platycheir*) and 2 (NGS-2: blast hit, Ischyroceridae sp.) in Table S3.

**S1 Table. Nanopore sequencing primers.**

**S2 Table. Detailed results of DNA barcoding for old nematode specimen using ONTbarcoder and NGSpeciesID.**

**S3 Table. Detailed results of DNA barcoding for isolated microorganisms using ONTbarcoder and NGSpeciesID.**

