## Supplemental Figure 1-3 for "Non-destructive DNA extraction from specimens and environmental samples using DESS preservation solution for DNA barcoding"

### Nematode specimen-5

```
consensus_2 TGTTTTAATTTTGCCCGGATTTGGAATTATTTCCCATATCACTCTTTATATGTCTGGTAA
consensus_3 TGTTTTAATTTTGCCCGGATTTGGAATTATTTCCCATATCACTCTTTATATGTCTGGTAA
            ************************************************************

consensus_2 AGATTCTATCTTTGGCAATATTGGTATAATTTATGCTATGTTGGTAATTGGAATTTTAGG
consensus_3 AGATTCTATCTTTGGCAATATTGGTATAATTTATGCTATGTTGGTAATTGGAATTTTAGG
            ************************************************************

consensus_2 TTGTGTGGTTTGAGCCCACCATATATTTACGGTTGGTTTAGATTTGGACACTCGAGCTTA
consensus_3 TTGTGTGGTTTGAGCCCACCATATATTTACGGTTGGTTTAGATTTGGACACTCGAGCTTA
            ************************************************************

consensus_2 TTTTACTTCAGCAACTATAATTATTGCTGTTCCTACAGGTGTAAAAATCTTTTCTTGATT
consensus_3 TTTTACTTCAGCAACTATAATTATTGCTGTTCCTACAGGTGTAAAAATCTTTTCTTGATT
            ************************************************************

consensus_2 GAGAACTTTATACGGTGGTAAAATAAAAATAAATATCCCTTTAATTTGGGCAATTGGTTT
consensus_3 GAGAACTTTATACGGTGGTAAAATAAAAATAAATATCCCTTTAATTTGGGCAATTGGTTT
            ************************************************************

consensus_2 TATTTTTTTATTTACTTTAGGGGGGTTAACAGGAATTGTTTTGTCTTCTTCCTCTTTAGA
consensus_3 TATTTTTTTATTTACTTTAGGGGG-TTAACAGGAATTGTTTTGTCTTCTTCCTCTTTAGA
            ************************ ***********************************

consensus_2 TGTTCTTCTCCATGATACTTATTATGTAGTT
consensus_3 TGTTCTTCTCCATGATACTTATTATGTAGTT
            *******************************
```

Figure S1

Sequence alignment showing the sequence similarity between consensus sequence 2 and 3 in specimen-5 (Table 4). Yellow boxed regions indicate the sequenced by Sanger sequencer and registered in NCBI.

### A Animal-1

```
consensus_1 ATATCTCCTCTCGCAGCAGCTACAGCTCATAGAGGTGCCTCTGTTGATCTAGCTATTTTT
consensus_2 -----TTTTTTAAGAGCAGCTATAGCTCATAGAGGAGGTGCAGTAGATTTAGCTATTTTT
                 *  * *   ******** ************ *   * ** *** ***********

consensus_1 -CTCTTCATTTAGCTGGAGCTTCCTCTATTCTCGGGGCTATTAATTTTATTTCTACAGTA
consensus_2 TCTCTTCATTTGGCAGGAGCTTCTTCTATTTTAGGAGCTATTAATTTTATTTCAACTGTA
             ********** ** ******** ****** * ** ***************** ** ***

consensus_1 ATTAATATACGAGCCCCTAATATAAAATTTGACCAAATACCTTTATTTGTTTGGTCCATT
consensus_2 ATTAATATGCGAACAGCAGGAATATTTATAGATCGTATACCTTTATTTGTTTGGTCTGTT
            ******** *** *  *    ***    * ** *  ********************  **

consensus_1 TTTATCACAGCCATTCTTTTACTTCTTTCTTTACCAGTTTTAGCTGGAGCCATTACTATA
consensus_2 TTTATTACTGCAATTTTATTATTACTTTCGTTACCTGTTTTAGCAGGAGCAATTACTATA
            ***** ** ** *** * *** * ***** ***** ******** ***** *********

consensus_1 TTATTAACAGACCGAAATTTAAATACTTCTTTTTTTGACCCTAGAGGAGGAGGAGACCCT
consensus_2 TTACTCACAGACCGTAATTTAAACACTTCATTTTTTGACCCATTAGGAGGAGGTGATCCT
            *** * ******** ******** ***** ***********   ********* ** ***

consensus_1 ATCCTTTACCAGCATCTTTTTT--
consensus_2 ATTCTTTATCAACATTTATTTTTA
            ** ***** ** *** * ****
```

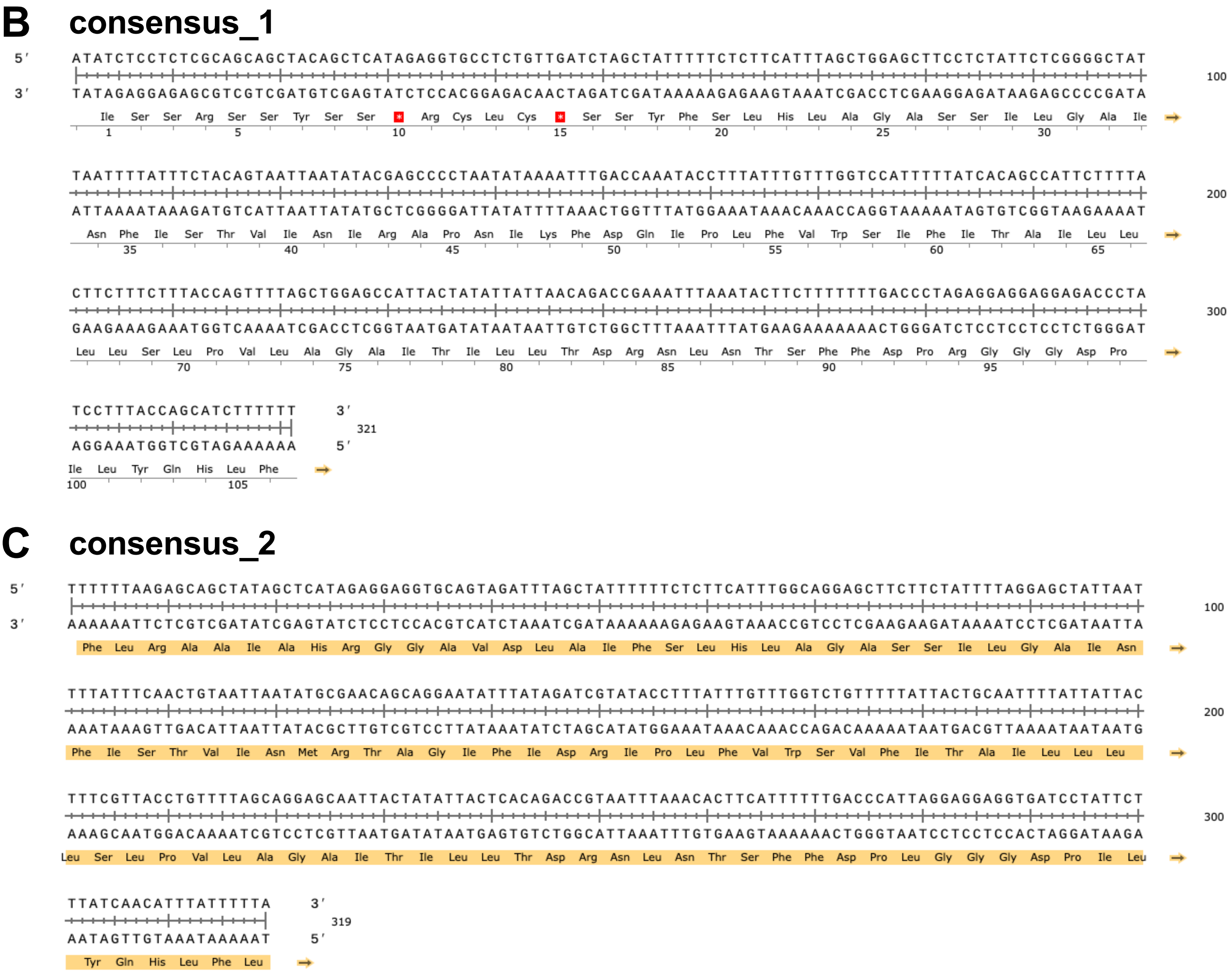


Figure S2

**A** Sequence alignment showing the sequence similarity between consensus sequence 1 (best hit: *Obesogammarus platycheir*) and 2 (best hit: Ischyroceridae sp.) in Animal-1 (Table 5). The sequence exhibit about 77% similarity (identity 250/324, gap 8/324). Red characters represent mismatch alignment. **B,C** Consensus sequence 1 (**B**) and 2 (**C**). The amino acid sequence was predicted. The asterisk (*) with a red background denotes the stop codon.

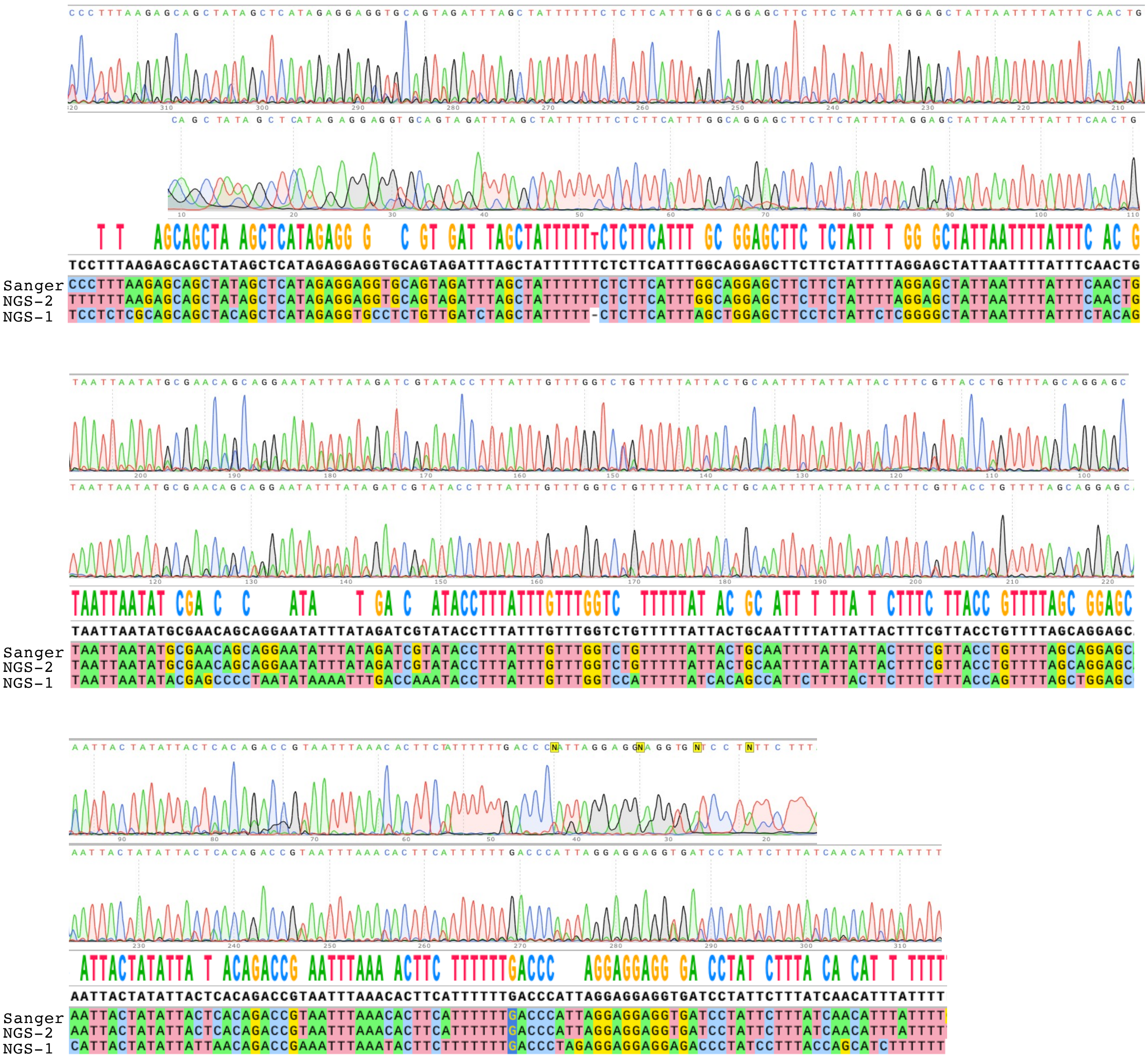


Figure S3

Sanger sequencing chromatograms of both forward and reverse strands for the COI region (mlCOIintf/HCO2198) in Animal-1 (Table 5). Sequence alignment showing the sequence similarity among Sanger and NGS outputs [consensus sequence 1 (NGS-1: blast hit, *Obesogammarus platycheir*) and 2 (NGS-2: blast hit, Ischyroceridae sp.) in Table S3].
